# DeepNeuropePred: a robust and universal tool to predict cleavage sites from neuropeptide precursors by protein language model

**DOI:** 10.1101/2023.07.07.547760

**Authors:** Lei Wang, Zilu Zeng, Zhidong Xue, Yan Wang

## Abstract

Neuropeptides play critical roles in many biological processes such as growth, learning, memory, metabolism, and neuronal differentiation. A few approaches have been reported for predicting neuropeptides that are cleaved from precursor protein sequences. However, these models for cleavage site prediction of precursors were developed using a limited number of neuropeptide precursor datasets and simple precursors representation models. In addition, a universal method for predicting neuropeptide cleavage sites that can be applied to all species is still lacking. In this paper, we proposed a novel deep learning method called DeepNeuropePred, using a combination of pretrained language model and Convolutional Neural Networks for feature extraction and predicting the neuropeptide cleavage sites from precursors. To demonstrate the model’s effectiveness and robustness, we evaluated the performance of DeepNeuropePred and four models from the NeuroPred server in the independent dataset and our model achieved the highest AUC score (0.916), which are 6.9%, 7.8%, 8.8%, and 10.9% higher than Mammalian (0.857), insects (0.850), Mollusc (0.842) and Motif (0.826), respectively. For the convenience of researchers, we provide an easy-to-install GitHub package (https://github.com/ISYSLAB-HUST/DeepNeuropePred) and a web server (http://isyslab.info/NeuroPepV2/deepNeuropePred.jsp).

**Key Points:**

1. DeepNeuropePred uses a deep learning algorithm based on protein language model to accurately predict neuropeptide cleavage sites from neuropeptide precursors.
2. Independent test experiments show that DeepNeuropePred achieves significantly better performance than existing methods.
3. DeepNeuropePred could capture meaningful patterns between neuro-peptide and non-neuropeptide cleavage sites.
4. We further provide an easy-to-install GitHub package and a web server.

## 1. Introduction

Neuropeptides are a diverse and complex class of signaling molecules that modulate nearly every physiological process and behavior in living species[1]. Typically consisting of less than 100 amino acids, they are produced from larger precursor molecules through a series of post-translational processing steps[2, 3]. Neuropeptides exert their effects not only via the nervous system but also peripherally through the endocrine system, where they regulate various functions, including food intake, metabolism, reproduction, fluid homeostasis, cardiovascular function, energy homeostasis, stress control, pain perception, social behaviors, memory and learning, and circadian rhythm[4-7]. Consequently, they are implicated in the pathogenesis of numerous diseases, and the neuropeptide signaling system represents a promising therapeutic target for the treatment of sleep disorders, autism, depression, heart failure, obesity, diabetes, high blood pressure, epilepsy, and other disorders[1, 4, 8, 9]. Furthermore, neuropeptides serve as valuable biomarkers and diagnostic probes for prospective disease diagnosis and prognosis. The precursor mRNA encodes a signal sequence and one or more neuropeptides, which can include multiple copies of the same neuropeptide. The sequence of prohormones can often be inferred from genetic information. However, it is generally difficult to predict biologically active peptides based on genetic information due to the many processing steps involved. Experimental validation of the final neuropeptide structure can also be challenging, as the neurons that synthesize specific neuropeptides are often rare, structurally complex, and present at low physiological concentrations[10].

With the development of the next-generation sequencing technology, more and more new genomes have been obtained and it is more important to identify precursors and detect neuropeptides. The successful combination of mass spectrometry with genome sequencing has enabled the characterization of the insect peptidome for neuropeptides in Drosophila melanogaster[11-13] and Apis mellifera[14]. While the most ubiquitous sites for cleavage are these basic residues, there are also some additional amino acid combinations that can be used as cleavage points in the prohormone[15]. Neuropeptide precursors contain certain structural information: (1) The signal peptide; (2) The basic motifs; (3) C-terminal amidation; (4) post-translational modifications[2]. Thus, due to the conserved pattern, it is possible to identify the cleavage sites of neuropeptide precursors. NeuroPred is a web application designed to predict cleavage sites at basic amino acid locations in neuropeptide precursor sequences, including Motif[10], Mammalian[16], Mollusc[17], and Insect[18] models. Southey et al. [10] proposed a Known Motif model comprised of several prevalent motifs associated with neuropeptide precursor cleavage. While the Motif method identified the most known cleavages, it also had a high rate of false positive prediction results. Hummon et al. [17] also predicted neuropeptide cleavage sites in mollusk precursors using the logistic regression model on combinations of amino acids and location information. Amare et al. [16] developed a binary logistic regression model to analyze mammalian neuropeptide precursors. The study revealed significant differences in processing between vertebrate and molluscan precursors, particularly in the processing of dibasic sites. NeuroPred-Insect model was trained on Apis mellifera and Drosophila melanogaster precursors using binary logistic regression, multi-layer perceptron and k-nearest neighbor models. All of the aforementioned neuropeptide cleavage site prediction tools are limited by small-scale data, resulting in unsatisfactory predictive accuracy.

In recent years, deep learning-based methods have been applied to different biological problems[19-22]. Compared with traditional machine learning models, deep learning-based methods can better capture the dataset distribution feature and reduce complex feature engineering processing. In the field of natural language processing (NLP), transfer learning through pre-trained language models has become ubiquitous. These models primarily learn context-based word embeddings, such as BERT[23]. With over a billion protein sequences in databases, a highly effective approach is to use self-supervised language models to learn latent information from unlabeled sequences. A pre-trained language model called ESM, based on the Transformer architecture, was trained to predict protein contact maps using over 680 million parameters[24]. Protein language models have presented some exciting breakthroughs, enabling the discovery of protein structure and function solely from the evolutionary relationships that exist in sequence corpora. Recently, different researchers[25-28] have developed some deep language models based on large-scale protein sequence datasets. These protein language models have been widely used in related tasks of the protein, such as signal peptide prediction[29], protein subcellular localization[30], bitter peptide[31], neuropeptide prediction[32], protein domain prediction[33, 34], and transmembrane protein prediction[35]. The use of transfer learning to obtain better feature representations for neuropeptide precursors is highly inspiring and holds great potential.

Currently, a universal and robust solution for predicting cleavage sites of neuropeptide precursors is still lacking. With more and more neuropeptide precursors being incorporated into protein public databases such as UniProt, this provides an opportunity for improved annotation of the cleavage site prediction of these neuropeptides. By collecting the neuropeptide precursors datasets from the UniProt database[36], we proposed a novel and robust model called DeepNeuropePred to detect the cleavage sites of neuropeptide precursors. Through a combination of pre-trained language models and Convolutional Neural Networks for feature extraction, our developed model achieved superior performance than four models from NeuroPred (Mammlian, Insects, Mollusc, and Motif). The DeepNeuropePred webserver is freely available at http://isyslab.info/NeuroPepV2/deepNeuropePred.jsp and source code is visible at https://github.com/ISYSLAB-HUST/DeepNeuropePred.

## 2. Materials and Methods

### 2.1. Benchmark Dataset

By searching the keywords such as “neuropeptide” from the UniProt/KB database and filtering these protein terms without the precursor of flags and the signal peptide, we collected 1194 complete reviewed precursors of neuropeptide. Here, we adopted 31 test precursors as the independent test dataset which was integrated into the UniProt database after 2014. To guarantee a fair comparison on the independent test dataset, the collected sequences of the training dataset that are similar to the test precursors at a threshold of 40% using CD-HIT[39] would be dropped. By the above steps, the remaining precursors from the training dataset included 717 precursors (training: validation=4:1). All training data and test data are freely available at https://github.com/isyslab-hust/DeepNeuropePred.

### 2.2. Feature extraction based on the protein language model

The emergence of pre-trained language models has propelled research on protein representation into an exciting new phase that eliminates the need for human annotation. This breakthrough allows for the acquisition of protein sequence representation from a vast pool of unannotated protein sequences, leading to significant improvements in downstream tasks. Additionally, utilizing pre-trained language models acts as an efficient regularization technique, reducing the risk of overfitting on limited training data.

In this context, we have adopted a transformer-based self-supervised language model called ESM, which boasts over 85 million parameters and 12 transformer layers. The ESM model is capable of processing protein sequences as inputs, generating dynamic embeddings with a dimension of *L* ∗ 768, where *L* represents protein length. The ESM model have been developed based on the scaled dot-product attention. The attention function is defined as:

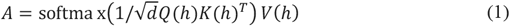

Here the query *Q*, key *K*, and value *V*, are projections of the protein sequence to *n* × *d* matrices where n is the length of the protein sequence and d is the dimension of the outer product between Q and K. This outer product obtains an *n* × *n* attention map, which is rescaled and passed through the SoftMax function, thereby representing each position of the sequence in the output as a convex combination of the sequence of values V. The pre-trained language model (ESM-12) can obtain the global feature representation of the precursor neuropeptides because the input is the full length of the precursor sequence rather than the window sequence of the cleavage site.

### 2.3. Cleavage site local information enhancement with convolutional neural network

The processing procedure of the complete precursor sequence was followed as the Insect model of NeuroPred (Southey, et al., 2008). Considering the strong local correlation of the neuropeptide cleavage site, we use a multi-scale convolutional neural network based on sparse connectivity to obtain the local vector representations of adjacent residues. The strong local correlation feature can be obtained by the followings:

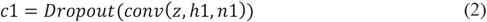

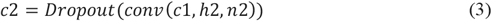

Where *conv* is the convolutional function, *h*1, *h*2 is the convolutional kernel size and *n*1, *n*2 is the number of convolutional kernels. The dropout function is a technique utilized to mitigate the risk of overfitting in models. It operates by randomly deactivating neurons and their corresponding connections, thereby preventing the network from relying too heavily on any one neuron. This strategy encourages all neurons to develop better generalization abilities.

### 2.4. Loss function

DeepNeuropePred takes the entire window sequence of L residues as input and outputs a probability score, where the score indicates whether the input site belongs to neuropeptide cleavage sites. The cross-entropy loss function is used as the optimization goal of the neural network. In our proposed model, the learning rate is 0.001, and the optimizer is Adam optimizer. The loss function is defined as:

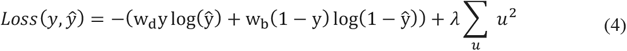

where *y* and *ŷ* are the label and the predicted possibility scores, respectively; ground truth label y reflects if it belongs to the neuropeptide cleavage site (1) or non-neuropeptide cleavage site (0), and *ŷ* is the output of the DeepNeuropePred; λ is the regularization factor; *u* represents all the trainable parameters; *w*_*d*_ and *w*_*b*_ are the reciprocal of the number of neuropeptide cleavage sites and non-neuropeptide cleavage sites, which can reduce the imbalance of samples in the dataset. Finally, 0.5 is selected as the threshold, probability scores greater than 0.5 is considered neuropeptide cleavage site, and those less than 0.5 is considered non-neuropeptide cleavage site.

### 2.5. Evaluation Metrics

To evaluate the model performance, we choose the following metrics: AUROC (Receiver Operating Characteristic curve), ACC (accuracy), Precision, Recall, AUPR (area under the precision–recall curve) and MCC (Matthew’s correlation coefficient). The relevant definitions are as follows:

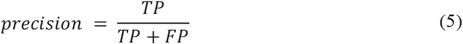

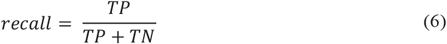

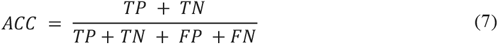

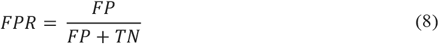

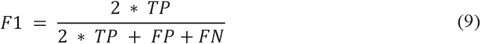

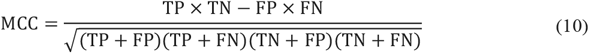

TP (true positive), FP (false positive), TN (true negative) and FN (false negative) are obtained from the confusion matrix. The ROC curve has the FPR (false positive rate) and TPR (recall or true positive rate) as the horizontal and vertical coordinates, respectively, and the area under the ROC curve is the AUC score. The area under the curve with recall and precision as the horizontal and vertical coordinates is the AUPR score.

## 3. Result and Discussion

### 3.1. Overview of the DeepNeuropePred framework

The model framework shown in Figure 1 consists of 4 parts, pre-trained self-supervised language model, convolutional layers, average pooling layer, and position-wise fully-connected layers. Using a pre-trained language model could reduce the risk of overfitting on small training data, which is equivalent to a kind of regularization method. The pre-trained language model (ESM-12, https://github.com/facebookresearch/esm) can obtain the global feature representation of the precursor because the input is the full length of the precursor sequence rather than the window sequence of the cleavage site. In addition, we integrated an advanced signal peptide prediction tool (SignalP5[37], https://services.healthtech.dtu.dk/services/SignalP-5.0/). Then, the processing procedure of the complete precursor sequence was followed as the Insect model of NeuroPred, and the candidate samples were fed into our model. Using the convolution layers with different scale kernels (1 and 3) could obtain local features of the windows of 18 amino acids in two different scales and the average pooling layer was used to obtain the glob representation of the windows. In the next stage, position-wise fully connected layers were used for mapping the embedding of cleavage sites and non-cleavage sites to the classification space.

**Figure 1.**
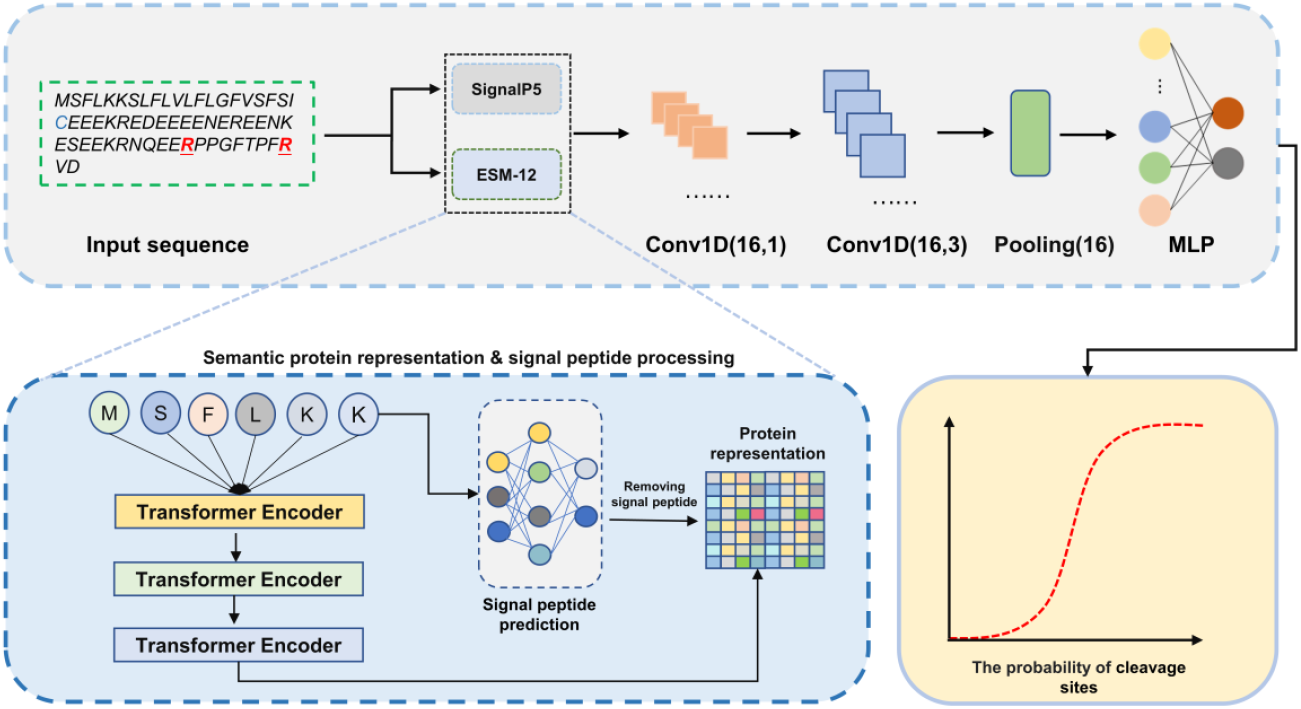
The flowchart of DeepNeuropePred.

**Figure 2.**
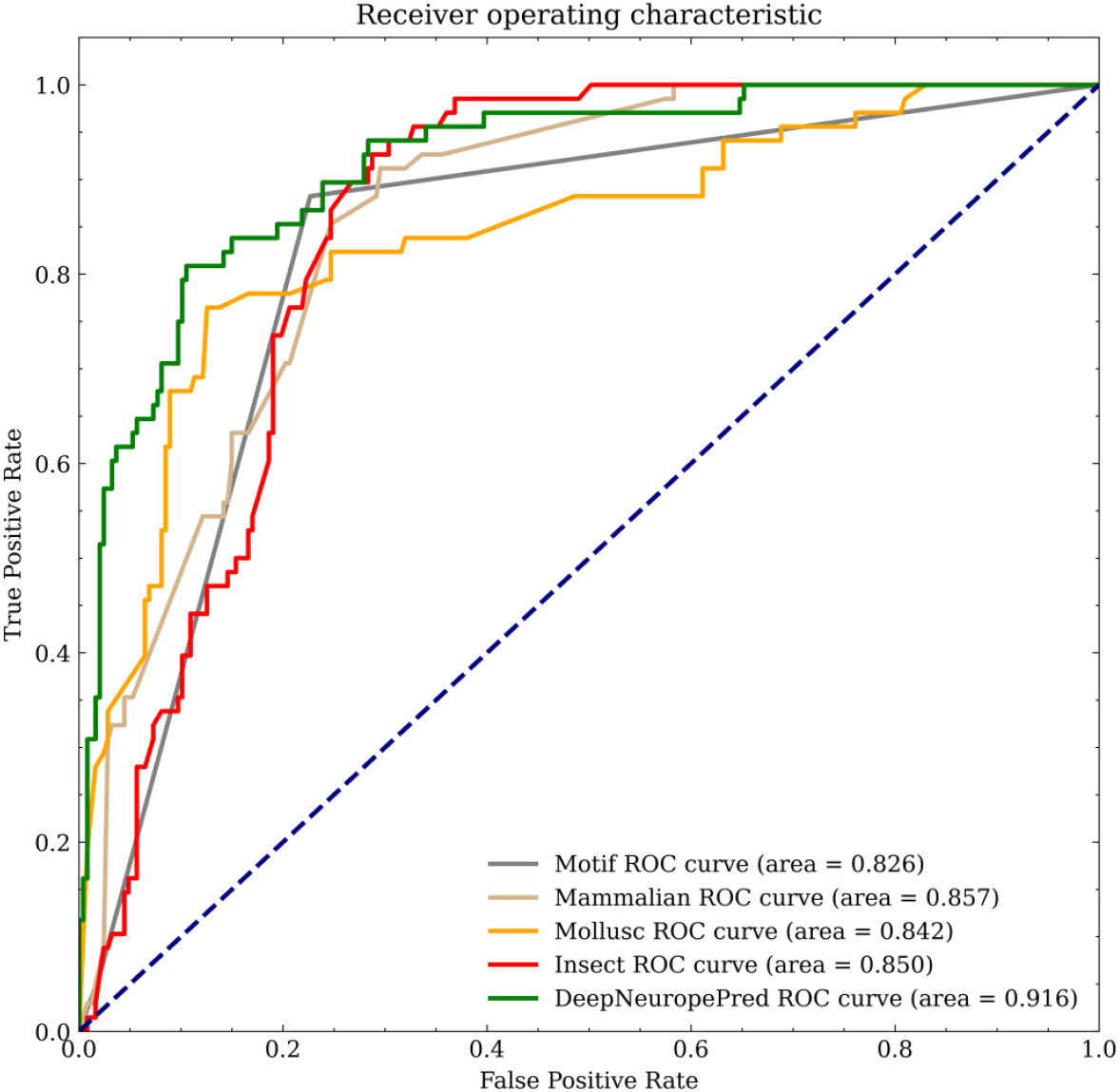
The Area Under the Receiver Operating Characteristic Curve of DeepNeuropePred, Motif, Mammalian, Mollusc, and Insect.

### 3.2. Ablation study of DeepNeuropePred

To evaluate the contribution of the CNN component (two convolutional layers) and ESM-12 module, we set up the ablative configurations. When the ESM-12 module was removed, the neuropeptide precursor sequences were encoded using one-hot encoding. When the two CNN layers were removed, a linear layer was used as a replacement. Five-fold cross-validation was used to evaluate the performance of DeepNeuropePred, DeepNeuropePred w/o ESM, and DeepNeuropePred w/o CNN on the training dataset. The entire training dataset was randomly split into five subsets containing the same number of chains. The results of the ablative experiment of neuropeptide cleavage site prediction are shown in Table 1.

**Table 1.** Five-fold cross-validation results of DeepNeuropePred, w/o ESM and w/o CNN.

| Model | Precision | Recall | F1-score | MCC | ACC | AUPRC |
| --- | --- | --- | --- | --- | --- | --- |
| w/o ESM | 0.79 | 0.83 | 0.81 | 0.63 | 0.81 | 0.72 |
| w/o CNN | 0.84 | 0.81 | 0.82 | 0.67 | 0.86 | 0.74 |
| DeepNeuropePred | **0.89** | **0.86** | **0.87** | **0.75** | **0.91** | **0.85** |

It is easy to see that the protein language model (ESM-12) obviously plays a very critical role in the neuropeptide cleavage site prediction. The ACC and MCC scores of DeepNeuropePred w/o ESM and DeepNeuropePred increased from 0.81 to 0.91, and 0.63 to 0.75, respectively. DeepNeuropePred achieved a Precision score of 0.89, a Recall score of 0.881, and an F1-score of 0.87 which was about 10% (0.79),3.0% (0.83), and 6.0% (0.81) higher than DeepNeuropePred w/o ESM. These results illustrate that the protein language module was also important for the neuropeptide cleavage site prediction. When the input part of ESM was removed, the performance of DeepNeuropePred degraded significantly. The ACC and MCC scores of DeepNeuropePred w/o CNN and DeepNeuropePred increased from 0.86 to 0.91, and 0.67 to 0.75, respectively. From this ablation experiment, it can be proved that the semantic representation based on the pre-trained model such as ESM can improve the prediction of neuropeptide cleavage sites. To make a fair and robust comparison, we further compared DeepNeuropePred with other state-of-the-art methods (four models from NeuroPred) on the independent test set.

### 3.3. Comparison with the state-of-the-art methods

We compared DeepNeuropePred with the existing models such as the Motif, Mammalian, Mollusc, and Insect models from the NeuroPred server. It should be emphasized that the test neuropeptide precursors are less 40% identity with the training dataset, which ensures the fairness of the comparison. As shown in Table 2, we see that DeepNeuropePred achieved the highest accuracy of 0.87, followed by Motif (0.80), Mammalian (0.78), Insect (0.78), and Mollusc (0.77). For the AUPRC score, DeepNeuropePred outperformed Motif by 30%, Mammalian by 22%, Insect by 29%, and Mollusc by 14%. Furthermore, DeepNeuropePred achieved the highest scores in Precision, Recall, MCC, and F1-score, indicating that it outperforms the four models from NeuroPred in predicting the cleavage sites of neuropeptides.

**Table 2.**
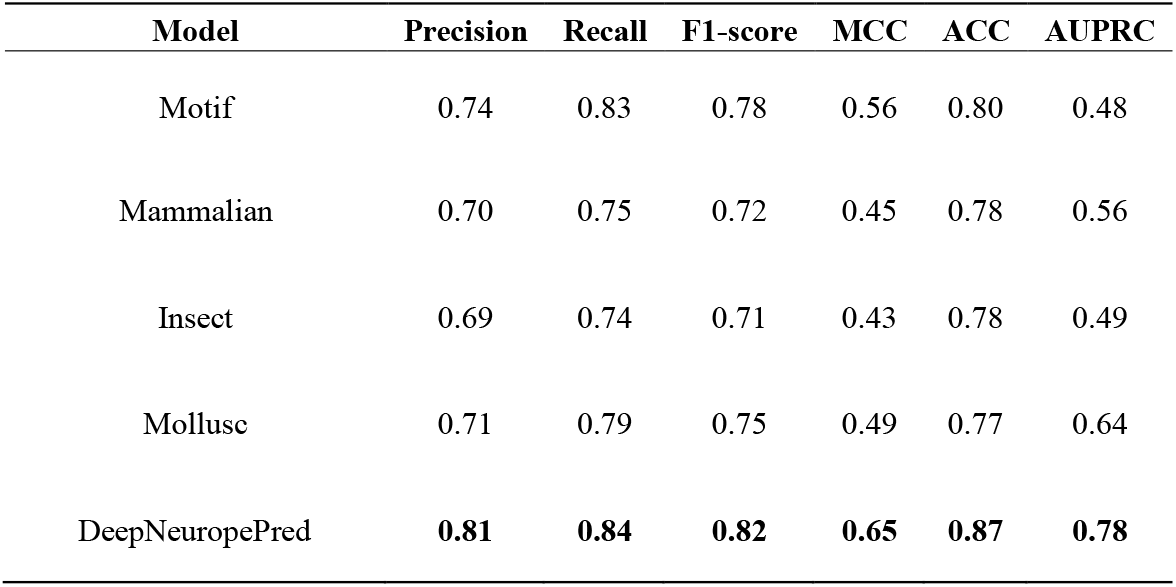
The performance metrics of DeepNeuropePred, Motif, Mammalian, Mollusc, and Insect.

In all test proteins, the positive samples of the cleavage site are 68 and the negative samples are 247(Positive: Negative=1:4) which is an unbalanced test set. However, the AUC score is insensitive to unbalanced data sets, and the performance of the model can be well evaluated. As shown in Figure2, DeepNeuropePred achieved the highest AUC score of 0.916, followed by Mammalian (0.857), Insect (0.850), Mollusc (0.842) and Motif (0.826), which are 6.9%, 7.8%, 8.8% and 10.9% higher respectively. Intuitively, the Motif model was based on some specific patterns, and the generalization ability is insufficient compared to other methods (Mammalian, Insect, and Mollusc). Furthermore, Mammalian, Insect, and Mollusc were trained by the specific domain datasets and these models always had a good performance for the similar precursors. However, our model proved its robustness through independent test precursors. These results also demonstrated the strong representation capability of DeepNeuropePred for both neuropeptide precursors and cleavage sites.

### 3.4. Visualization of features extracted by DeepNeuropePred

DeepNeuropePred is expected to capture meaningful patterns between neuro-peptide and non-neuropeptide cleavage sites. To investigate whether the DeepNeuropePred model has learned to encode cleavage site classifying attributes in its feature representation, we used all datasets of cleavage sites and non-cleavage sites and projected the learned embeddings of the ESM-12, convolutional layers 1 and 2 using t-distributed stochastic neighbor embedding (t-SNE) algorithm[38], which was decomposed into two dimensions. The parameters for t-SNE were set to a perplexity of 10 and 1000 iterations for the optimization. The results, as shown in Figure 3, clearly indicate that the embedding vectors of the Transformer blocks are unable to distinguish between cleavage sites and non-cleavage sites. Different convolutional layers enhance the inter-class distance between them, which indicates that local information enhancement is important for predicting cleavage sites. It can be concluded from Figure 3 that different layers can encode and capture features at different levels from the embedding feature of ESM-12.

**Figure 3.**
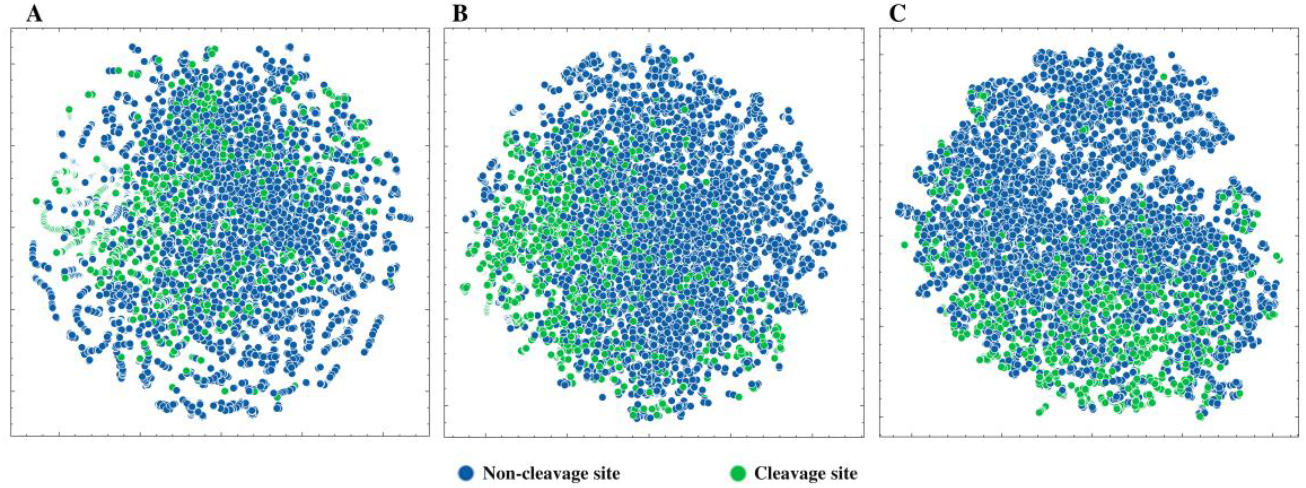
The neuropeptide cleavage sites are represented in the output embeddings of the ESM-12 (A), convolutional layer 1(B), and convolutional layer 2(C), visualized here with t-SNE.

### 3.5. Webserver of DeepNeuropePred

For the convenience of academic users, the DeepNeuropePred server is freely available at http://isyslab.info/NeuroPepV2/deepNeuropePred.jsp. Our web services are mainly built with Flask, Redis, and Celery. Flask (https://github.com/pallets/flask) and Celery (https://github.com/celery/celery) are two widely-used Python libraries that offer rich functionalities and APIs to help developers easily build complex asynchronous applications. The DeepNeuropePred webserver architecture is shown in Figure 4. When using Flask and Celery to build asynchronous tasks, we first create a Flask backend program and configure Celery’s message broker and result backend within the application. We then define a Celery task that takes two parameters and simulates the execution of a neural peptide cleavage site prediction task, with the Celery worker listening to the task queue. The Flask application is started and asynchronously calls the Celery task, immediately returning the task ID to the frontend. The task execution process occurs in the background, asynchronously executed by the Celery worker. Once the task is completed, the Celery worker stores the result in the result backend, which can be obtained by a frontend request. Throughout the process, Flask and Celery work hand-in-hand, with Flask being responsible for receiving client requests and asynchronously calling the Celery task, while Celery executes the task and stores the result in the result backend. This approach improves the response speed and performance of our application while ensuring the reliability and stability of tasks. For the DeepNeuropePred inference, the PyTorch framework (https://pytorch.org/) is used to construct the neural network.

**Figure 4.**
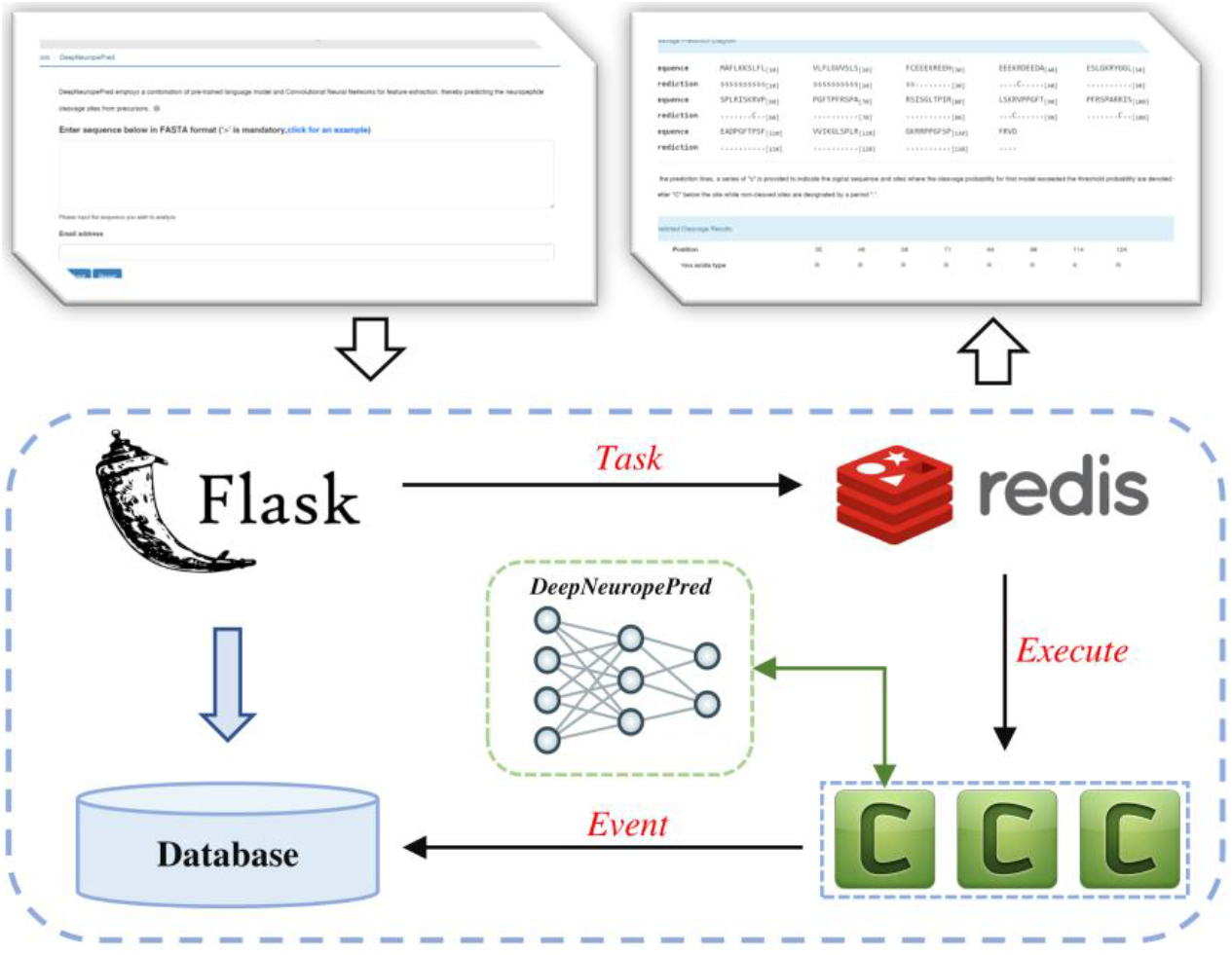
The front-end and back-end architecture of the DeepNeuropePred webserver.

## 4. Conclusion

In this study, we introduce DeepNeuropePred, a transfer learning method for detecting cleavage sites in neuropeptide precursors. Our approach was tested on a more rigorous dataset and offers the distinct advantage of being able to model the long-distance representation of the entire sequence, as opposed to just the local window feature of the cleavage site. In independent tests, DeepNeuropePred outperformed the existing model from the NeuroPred server. To the best of our knowledge, this is the first application of a transfer learning algorithm to neuropeptide cleavage site prediction. The use of transfer learning offers new possibilities for predicting cleavage sites in neuropeptide precursors with limited training datasets. In the future, we aim to further improve the performance of our method by developing new deep-learning techniques.

## Author Contributions

Conceptualization, L.W., Z.X. and Y.W.; Data curation, L.W. and Y.W.; Formal analysis, L.W., Z.X., Z.Z. and Y.W.; Funding acquisition, L.W., Z.X. and Y.W.; Methodology, L.W. and Y.W.; Project administration, Z.X. and Y.W.; Software, L.W.; Writing – original draft, L.W., Z.X., Z.Z. and Y.W.; Writing – review & editing, L.W., Z.X., Z.Z. and Y.W. All authors have read and agreed to the published version of the manuscript.

## Acknowledgments

Thanks to the Facebook Research team for providing the pre-trained weights for the transformer protein language models.

## References

1. Mendel HC, Kaas Q, Muttenthaler M. Neuropeptide signalling systems - An underexplored target for venom drug discovery, Biochem Pharmacol 2020;181:114129.

2. Burbach JP. What are neuropeptides?, Methods Mol Biol 2011;789:1–36.

3. Wang Y, Wang M, Yin S et al. NeuroPep: a comprehensive resource of neuropeptides, Database (Oxford) 2015;2015:bav038.

4. Hokfelt T, Broberger C, Xu ZQ et al. Neuropeptides--an overview, Neuropharmacology 2000;39:1337–1356.

5. Sobrino Crespo C, Perianes Cachero A, Puebla Jimenez L et al. Peptides and food intake, Front Endocrinol (Lausanne) 2014;5:58.

6. Shahjahan M, Kitahashi T, Parhar IS. Central pathways integrating metabolism and reproduction in teleosts, Front Endocrinol (Lausanne) 2014;5:36.

7. Kormos V, Gaszner B. Role of neuropeptides in anxiety, stress, and depression: from animals to humans, Neuropeptides 2013;47:401–419.

8. Nassel DR, Zandawala M. Recent advances in neuropeptide signaling in Drosophila, from genes to physiology and behavior, Prog Neurobiol 2019;179:101607.

9. Nassel DR. Neuropeptides in the nervous system of Drosophila and other insects: multiple roles as neuromodulators and neurohormones, Prog Neurobiol 2002;68:1–84.

10. Southey BR, Rodriguez-Zas SL, Sweedler JV. Prediction of neuropeptide prohormone cleavages with application to RFamides, Peptides 2006;27:1087–1098.

11. Baggerman G, Cerstiaens A, De Loof A et al. Peptidomics of the larval Drosophila melanogaster central nervous system, J Biol Chem 2002;277:40368–40374.

12. Baggerman G, Boonen K, Verleyen P et al. Peptidomic analysis of the larval Drosophila melanogaster central nervous system by two-dimensional capillary liquid chromatography quadrupole time-of-flight mass spectrometry, J Mass Spectrom 2005;40:250–260.

13. Predel R, Wegener C, Russell WK et al. Peptidomics of CNS-associated neurohemal systems of adult Drosophila melanogaster: a mass spectrometric survey of peptides from individual flies, J Comp Neurol 2004;474:379–392.

14. Hummon AB, Amare A, Sweedler JV. Discovering new invertebrate neuropeptides using mass spectrometry, Mass Spectrom Rev 2006;25:77–98.

15. Hummon AB, Huang HQ, Kelley WP et al. A novel prohormone processing site in Aplysia californica: the Leu-Leu rule, Journal of Neurochemistry 2002;82:1398–1405.

16. Amare A, Hummon AB, Southey BR et al. Bridging neuropeptidomics and genomics with bioinformatics: Prediction of mammalian neuropeptide prohormone processing, J Proteome Res 2006;5:1162–1167.

17. Hummon AB, Hummon NP, Corbin RW et al. From precursor to final peptides: A statistical sequence-based approach to predicting prohormone processing, Journal of Proteome Research 2003;2:650–656.

18. Southey BR, Sweedler JV, Rodriguez-Zas SL. Prediction of neuropeptide cleavage sites in insects, Bioinformatics 2008;24:815–825.

19. Shi Q, Chen W, Huang S et al. Deep learning for mining protein data, Brief Bioinform 2021;22:194 –218.

20. He Y, Shen Z, Zhang Q et al. A survey on deep learning in DNA/RNA motif mining, Brief Bioinform 2021;22.

21. Xu J, Li F, Leier A et al. Comprehensive assessment of machine learning-based methods for predicting antimicrobial peptides, Briefings in Bioinformatics 2021;22:bbab083.

22. Shi Q, Chen W, Huang S et al. DNN-Dom: predicting protein domain boundary from sequence alone by deep neural network, Bioinformatics 2019;35:5128–5136.

23. Devlin J, Chang M-W, Lee K et al. BERT: Pre-training of Deep Bidirectional Transformers for Language Understanding. 2019, 4171–4186.

24. Rives A, Meier J, Sercu T et al. Biological structure and function emerge from scaling unsupervised learning to 250 million protein sequences, Proc Natl Acad Sci U S A 2021;118.

25. Elnaggar A, Heinzinger M, Dallago C et al. ProtTrans: Toward Understanding the Language of Life Through Self-Supervised Learning, IEEE Transactions on Pattern Analysis and Machine Intelligence 2022;44:7112–7127.

26. Geffen Y, Ofran Y, Unger R. DistilProtBert: a distilled protein language model used to distinguish between real proteins and their randomly shuffled counterparts, Bioinformatics 2022;38:ii95–ii98.

27. Alley EC, Khimulya G, Biswas S et al. Unified rational protein engineering with sequence-based deep representation learning, Nat Methods 2019;16:1315–1322.

28. Bepler T, Berger B. Learning the protein language: Evolution, structure, and function, Cell Syst 2021;12:654–669 e653.

29. Teufel F, Armenteros JJA, Johansen AR et al. SignalP 6.0 predicts all five types of signal peptides using protein language models, Nature Biotechnology 2022;40:1023-+.

30. Thumuluri V, Almagro Armenteros JJ, Johansen AR et al. DeepLoc 2.0: multi-label subcellular localization prediction using protein language models, Nucleic Acids Res 2022.

31. Jiang J, Lin X, Jiang Y et al. Identify Bitter Peptides by Using Deep Representation Learning Features, Int J Mol Sci 2022;23.

32. Wang L, Huang C, Wang M et al. NeuroPred-PLM: an interpretable and robust model for neuropeptide prediction by protein language model, Brief Bioinform 2023;24.

33. Wang L, Zhong H, Xue Z et al. Res-Dom: predicting protein domain boundary from sequence using deep residual network and Bi-LSTM, Bioinform Adv 2022;2:vbac060.

34. Wang L, Wang Y. GNN-Dom: An Unsupervised Method for Protein Domain Partition via Protein Contact Map. In: Bioinformatics Research and Applications: 18th International Symposium, ISBRA 2022, Haifa, Israel, November 14–17, 2022, Proceedings. 2023, p. 286–294. Springer.

35. Wang L, Zhong HL, Xue ZD et al. Improving the topology prediction of a-helical transmembrane proteins with deep transfer learning, Computational and Structural Biotechnology Journal 2022;20:1993–2000.

36. UniProt C. UniProt: the universal protein knowledgebase in 2021, Nucleic Acids Res 2021;49:D480–D489.

37. Almagro Armenteros JJ, Tsirigos KD, Sonderby CK et al. SignalP 5.0 improves signal peptide predictions using deep neural networks, Nat Biotechnol 2019;37:420–423.

38. Van der Maaten L, Hinton G. Visualizing data using t-SNE, Journal of Machine Learning Research 2008;9.

39. Fu L, Niu B, Zhu Z et al. CD-HIT: accelerated for clustering the next-generation sequencing data, Bioinformatics 2012;28:3150–3152.

